# Enhanced meningeal lymphatic drainage ameliorates neuroinflammation and hepatic encephalopathy in cirrhotic rats

**DOI:** 10.1101/2020.05.01.072611

**Authors:** Shao-Jung Hsu, Chihao Zhang, Jain Jeong, Seong-il Lee, Matthew McConnell, Teruo Utsumi, Yasuko Iwakiri

## Abstract

**Background and aims:** Hepatic encephalopathy (HE) is a serious neurological complication in patients with liver cirrhosis. Nothing is known about the role of the meningeal lymphatic system in HE. We tested our hypothesis that enhancement of meningeal lymphatic drainage could decrease neuroinflammation and ameliorate HE.

**Methods:** A 4-week bile duct ligation (BDL) model was used to develop cirrhosis with HE in rats. Brain inflammation in patients with HE was evaluated using archived GSE41919. Motor function of rats was assessed by the rotarod test. AAV8-VEGF-C was injected into the cisterna magna of BDL rats one day after surgery to induce meningeal lymphangiogenesis.

**Results:** Cirrhotic rats with HE showed significantly increased microglia activation in the middle region of the cortex (*p*<0.001) as well as increased neuroinflammation as indicated by significant increases in IL-1β, INFγ, TNFα and Iba1 expression in at least one of the three regions of the cortex. Motor function was also impaired in rats with HE (*p*<0.05). Human brains of cirrhotic patients with HE also exhibited upregulation of pro-inflammatory genes (NF-κβ, Iba1, TNFα and IL-1β) (n=6). AAV8-VEGF-C injection significantly increased meningeal lymphangiogenesis (*p*=0.035) and tracer dye uptake in the anterior and middle regions of the cortex (*p*=0.006 & 0.003, respectively), their corresponding meninges (*p*=0.086 & 0.006, respectively) and the draining lymph nodes (*p*=0.02). Further, AAV8-VEGF-C decreased microglia activation (*p*<0.001) and neuroinflammation, and ameliorated motor dysfunction (*p*=0.024).

**Conclusion:** Promoting meningeal lymphatic drainage and enhancing waste clearance improves HE. Manipulation of meningeal lymphangiogenesis could be a new therapeutic strategy for the treatment of HE.

## Introduction

Hepatic encephalopathy (HE) is a disabling neurological complication of cirrhosis which causes cognitive, psychiatric, and motor impairment. In the United States, HE drives large numbers of cirrhosis-related hospitalizations and places a tremendous burden on the healthcare system^1, 2^. A key pathogenic factor for HE in cirrhotic patients is believed to be high levels of ammonia in the circulation. This hyperammonemia results from a decreased detoxification capacity in the liver as a result of liver failure and/or the development of porto-systemic shunts that allow blood flow to bypass the liver^3, 4^. Therefore, lowering blood ammonia levels is an important measure to ameliorate HE^5^ and is the target of our current treatments. However, elevated ammonia is not the sole pathogenic factor in HE, because patients may have HE with normal ammonia levels. Neuroinflammation, acting in synergy with ammonia, is increasingly recognized as a key pathogenic factor in HE and is a critical target for a new therapeutic approach to this debilitating condition^6^.

Because it is a highly metabolic organ, the brain needs an efficient drainage system to clear waste materials. While in peripheral tissues lymphatic vessels help to drain excess fluid and macromolecules, the central nervous system (CNS: the brain and spinal cord) was long thought to lack lymphatic drainage. However, the recent discoveries of the unique clearance and drainage systems in the brain, i.e., the glymphatic (glial-lymphatic) system^7^ and the meningeal lymphatic system^8, 9^, have dramatically advanced our understanding of how toxic materials can be removed from the brain. In the brain, the glymphatic system helps to direct cerebral spinal fluid (CSF) from the periarterial space to the brain parenchyma, where CSF is mixed with interstitial fluid containing cerebral waste products, such as metabolites and proteins^7^. CSF then enters into the perivenous space through the brain parenchyma and drains to the subarachnoid space. A recent study by Jalan and colleagues demonstrated that glymphatic function was altered in cirrhotic rats with HE, suggesting potential involvement of the glymphatic system in the development of HE^10^.

Meningeal lymphatic vessels, located underneath the skull, were discovered in 2015^8, 9^ and found to be the major route for discharging brain waste materials and for trafficking immune cells from the brain. CSF filled with waste materials and immune cells in the subarachnoid space enters into meningeal lymphatic vessels and drains to collecting lymphatic vessels outside the brain and then to lymph nodes in the neck called the deep cervical lymph nodes (dCLN)^8, 9, 11–14^. Because of these functions, the meningeal lymphatic system has received tremendous attention as a potential therapeutic target for treating neurodegenerative and neuroinflammatory disorders. Currently, the role of the meningeal lymphatic system in HE remains to be explored.

Several studies of rats with HE have shown the presence of neuroinflammation in the brain^15, 16^. Neuroinflammation is caused by various mediator molecules, such as cytokines, chemokines and reactive oxygen species, which are produced by various cell types, including microglia, astrocytes, endothelial cells, and peripherally derived immune cells. Microglia are the brain’s resident macrophages and play a central role in its innate immune capacity. Chronic brain inflammation due to activated microglia and increased production of pro-inflammatory cytokines, such as IL-1β, TNFα and IFNγ, could be related to pathological features of HE such as edema and increased brood-brain barrier (BBB) permeability^17^.

We hypothesized that enhanced meningeal lymphatic drainage might reduce microglial activation and brain inflammation and thereby ameliorate HE in cirrhotic rats. VEGF-C is the most potent lymphangiogenic factor, which binds to its receptor VEGFR3 (FLT4) and induces proliferation and migration of lymphatic endothelial cells, thereby leading to lymphangiogenesis (i.e., new lymphatic vessel formation)^18^. We found that AAV-VEGF-C delivery through the cisterna magna led to increased VEGF-C and lymphangiogenesis in the meninges as well as increased meningeal lymphatic drainage to the dCLN in cirrhotic rats. Further, AAV-VEGF-C decreased microglia activation and inflammation in the brain and improved motor dysfunction resulting from HE.

## Materials and methods

### Animal model: bile duct ligation

A 4-week bile duct ligation (BDL) model was used to develop hepatic encephalopathy (HE) in rats as shown in other studies^10^. Male Sprague-Dawley rats approximately 2 months of age (average weight: 320.5 g) were subjected to sham or common BDL surgery. In brief, under anesthesia with ketamine (75 mg/kg) and xylazine (5 mg/kg), the common bile duct was exposed and ligated with 3-0 silk sutures at two locations. The first ligature was made below the junction of the hepatic ducts and the second ligature above the entrance of the pancreatic duct, followed by resection of the common bile duct between the ligatures. To avoid coagulation defects, all rats, including sham rats for control, received weekly vitamin K injection (50 μg/kg)^19^. All animal experiments were approved by the Institutional Animal Care and Use Committee of the Veterans Affairs Connecticut Healthcare System and performed in adherence with the National Institutes of Health Guide for the Care and Use of Laboratory Animals.

### Intra-cisterna magna tracer injection

A 66 kDa fluorescent tracer, ovalbumin conjugated with Alexa Flour 647 (OVA-647, Thermo Fisher Scientific, Waltham, MA), was injected into the cisterna magna of rats under anesthesia with ketamine (75 mg/kg) and xylazine (5 mg/kg) to evaluate glymphatic function and meningeal lymphatic drainage. Depth of anesthesia was kept consistent among rats to avoid a differential influence of anesthesia on their glymphatic function^20^. An anesthesia protocol was adapted from a previous study with slight modifications^21^. Depth of anesthesia was monitored by the pedal reflex and respiratory rate. An additional 1/20 of the initial dose was administered when a rat responded to a toe pinch or had over 30 breaths per minute. After stabilization, a further boost of ketamine and xylazine was prohibited. Only those rats with respiratory rates between 20 and 30 breaths per minute during tracer injection were used.

A 30-gauge needle connected with a PE10 tube filled with the tracer (0.5 mg/ml in artificial cerebrospinal fluid) was inserted 3 mm deep into the cisterna magna. Glue was attached to the insertion site. 80 μl of the tracer was administered at a rate of 1.6 μl/min by a microliter syringe (Hamilton Company, Reno, NV) using a syringe pump (Harvard Apparatus, Holliston, MA). This flow rate was shown to have no effects on intracranial pressure^10, 22^. The injection was stopped in 50 minutes and the needle was removed 10 minutes after completion of injection to prevent backflow of the tracer. Rats were then sacrificed to collect blood and tissues including the brain, skull, and deep cervical lymph nodes (dCLN).

### Intra-cisterna magna AAV8-VEGF-C injection

One day after BDL surgery, rats were randomly divided into two groups, one given adeno-associated virus (AAV)8-mouseVEGF-C (Vector Biolabs, Malvern, PA) and the other given AAV8-GFP (control) (Addgene, Watertown, MA), to examine the effect of VEGF-C overexpression on the brain lymphatic system and HE. These AAVs were prepared in phosphate-buffered saline (PBS) with a concentration of 1.5 x 10^11^ GC/rat in a total volume of 20 μl on the same day of injection.

Similar to the tracer injection above, a 30-gauge needle connected with a PE10 tube filled with AAV solution was inserted 3 mm deep into the cisterna magna under anesthesia with ketamine (75 mg/kg) and xylazine (5 mg/kg). A 20 μl of solution was injected into the cisterna magna at a rate of 1.6 μl/min (1.5 x 10^11^ GC/rat in both groups) by a microliter syringe (Hamilton Company) using a syringe pump (Harvard Apparatus). The needle was removed 10 minutes after completion of injection and the wound was closed. All samples were collected 4 weeks after BDL surgery.

### Tissue preparation for evaluation of brain glymphatic function and meningeal lymphatic *drainage*

(Refer to the Supplemental materials section)

### Immunofluorescence analysis of the brain and meninges

(Refer to the Supplemental materials section)

### Quantification of meningeal lymphangiogenesis

(Refer to the Supplemental materials section)

### Quantification of microglia activation

A quantification method for microglia activation was adapted from a previous study with slight modifications^23^. Images were obtained using a fluorescent microscope (AX10, Zeiss) with a 40x objective and blindly analyzed by ImageJ with the AnalyzeSkeleton (2D/3D) plugin. First, an image was sharply contrasted by unsharp masking (settings: pixel radius of 3 and mask weight of 0.8). Then, salt-and-pepper noise generated by unsharp masking was removed by the Despeckle. The image was converted to a binary image by threshold function. Close and despeckle functions were applied to the image to make the contour of microglia more clear. Using the paintbrush tool, cells of interest were chosen. Fragmented processes were connected by matching photomicrograph as a reference. Then, the image was skeletonized and analyzed with the analyze skeleton function. Microglia activation was assessed by endpoint numbers and total branch lengths per cell. In each sample, 10 images were analyzed with each image including 1 to 4 cells.

### Measurement of key metabolites

(Refer to the Supplemental materials section)

### Quantitative real-time PCR

Total RNA was isolated from the anterior, middle and posterior regions of the cerebral cortex as well as the whole meninges. While still frozen, these samples were ground to powder using a mortar and a pestle in the presence of liquid nitrogen. Total RNA was isolated from powdered samples using TRIzol (QIAGEN, Germantown, MD) according to the manufacturer’s instructions. One microgram of the total RNA was reverse-transcribed using the Transcriptor First Strand cDNA Synthesis Kit (Roche Diagnostics, Florham Park, NJ). qPCR was performed using iTaq Universal SYBR Green Supermix (Bio-Rad Laboratories) and analyzed by the 7500 Real-Time PCR System (Applied Biosystems, Foster City, CA). All oligonucleotide primer sets for qPCR are presented in Supplemental Table 1. The oligonucleotide primers were all custom designed and synthesized by the Keck Oligonucleotide Synthesis facility at Yale School of Medicine. Relative mRNA expression levels were calculated based on the *2*^-ΔΔ*CT*^ *method and normalized to the 18S ribosomal RNA gene*.

### Rotarod performance test

Rotamex-5 (Columbus Instruments, Columbus, OH) was used for testing the ability of rats to stay on a rotating rod^24^.

(Refer to the Supplemental materials section for details)

### RNA-seq library preparation and sequencing

(Refer to the Supplemental materials section)

### RNA sequencing analysis of the middle region of the cortex in rat brains

Raw fastq files were assessed for quality before and after trimming adaptors using the *FastQC* software. Using *Trimmomatic* 0.36, reads with a minimum length of 50 bp and a Phred quality score of more than 30 were finally retained. The reference genome for rats was downloaded from the database available at the University of California, Santa Cruz (https://genome.ucsc.edu/cgi-bin/hgTables) and the latest version (RGSC 6.0/rn6) was chosen for assembly. We constructed indexes for a genome file and aligned the reads to the reference genome using *HISAT2*. The number of fragments mapped to each gene was calculated using the *featureCounts* command and used for statistical analysis.

Differential gene analysis was performed between samples by the *DESeq2* package. Raw data was normalized using the *trimmed mean of means (TMM)* algorithm built in *DESeq2*. Genes with a fold change of more than 1.5 or less than 2/3 as well as a *p* value of less than 0.05 were considered differentially expressed. To illustrate gene expression changes among different conditions, a venn diagram and heatmaps are presented in Figure 6 and Supplemental Figure 5. Gene Set Enrichment Analysis (GESA) was conducted to identify potential mechanisms by which VEGF-C could alleviate hepatic encephalopathy. We set 1000-time permutations for all pathways. A pathway was considered significantly enriched when a *p* value and a false discovery rate (FDR) were less than 0.05 and 0.25, respectively.

### Transcriptome analysis of human brains

An mRNA expression profile (GSE41919) in the study by Görg B et al.^25^ was downloaded from the Gene Expression Omnibus database (GEO, https://www.ncbi.nlm.nih.gov/geo/). This dataset consists of 8 brain samples from cirrhotic patients with hepatic encephalopathy and 8 brain samples from non-cirrhotic controls (platform: GPL14550, Aglient-028004 SurePrint G3 Human GE 8×60K Microarray). We annotated gene symbols based on the above platform. Expression values were averaged when duplicate data were presented. To assess the quality of the samples, hierarchical clustering and principal component analysis (PCA) were carried out using *hclust* and *FactoMineR* packages, respectively. When a particular sample from one group clustered with samples from the other group, it meant that the quality of that sample was not high enough to be included for the subsequent analysis.

To examine differentially expressed genes (DEGs), the *limma* package was used with an adjusted *p*-value of less than 0.05. Further, |log_2_ fold change (FC)|≥1 was employed as a threshold for identifying DEGs. For visualization, a heatmap was drawn for the top 30 DEGs using the *pheatmap* package. In addition, a volcano plot was generated to show a comprehensive picture of differential genes using the *ggplot2* package in the R language. Our workflow for the bioinformatic analysis of the GSE41919 dataset is shown in Figure 2A.

### Statistical analysis

Results are expressed as mean ± SEM. Student’s t test was used for statistical analysis, except for qPCR results in Figure 1B, in which Welch’s t-test was used because two groups (sham and BDL) displayed different variances. A *p* value of less than 0.05 was considered statistically significant.

**Figure 1.**
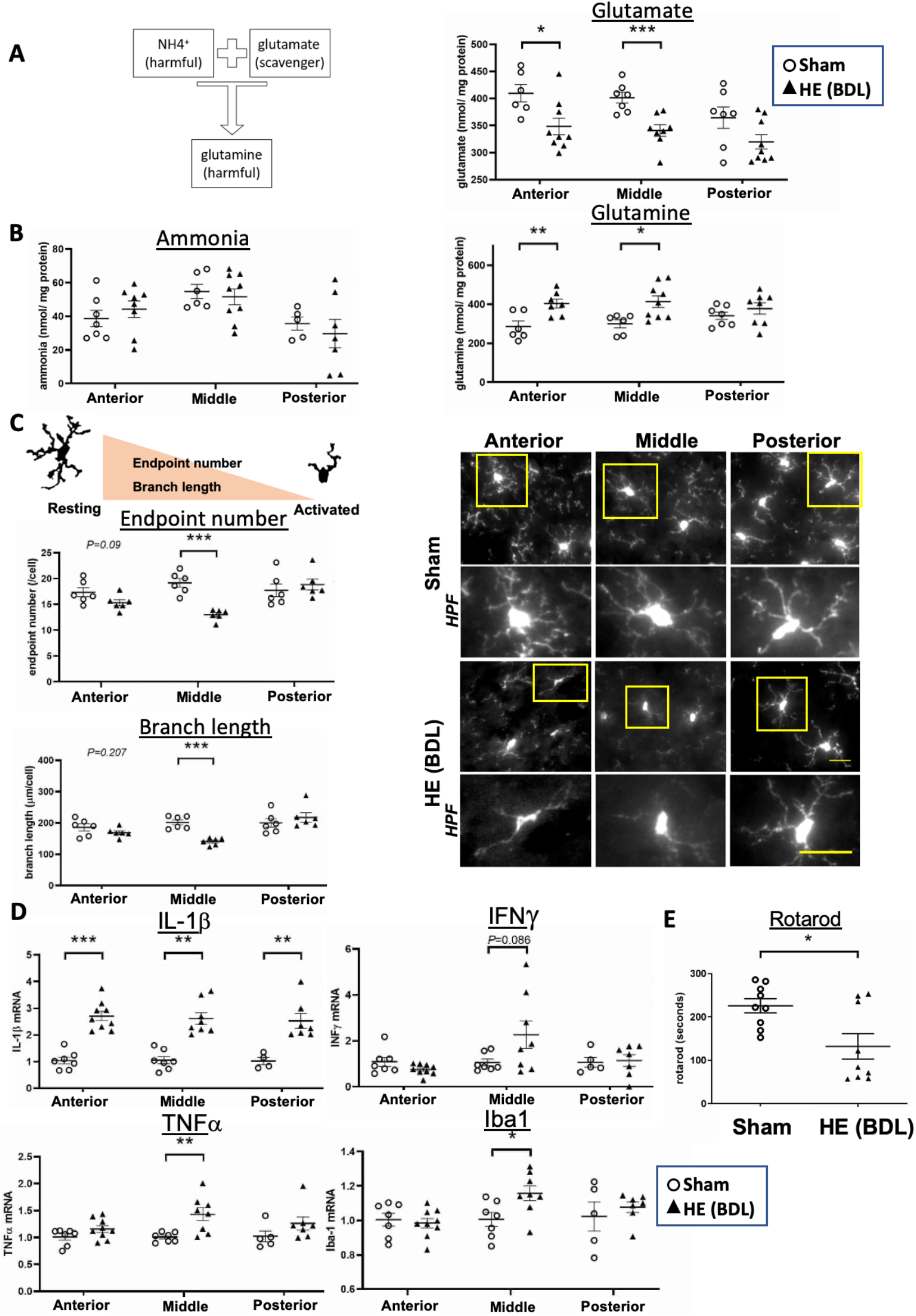
Increased brain inflammation in the anterior and middle regions of the cortex causes impaired motor function in BDL rats. **(A)** In hepatic encephalopathy, systemic hyperammonemia increases ammonia levels in the brain. Ammonia can be scavenged by glutamate, but as a result, toxic glutamine is formed and may contribute to the progression of hepatic encephalopathy. **(B)** Ammonia, glutamate and glutamine levels in the anterior, middle and posterior regions of the cortex in sham and BDL rats. n=4-7 (sham) and 7-9 (BDL). **(C)** Microglia activation was determined by the endpoint number and branch length of each cell. Activated microglia exhibit decreased endpoint numbers and branch lengths. Microglia were immunolabeled with Iba1 and examined for their activation in the anterior, middle and posterior regions of the cortex. HPF: high power field of a corresponding yellow square. Scale bar: 25μm. n=6 per group. **(D)** mRNA expression of pro-inflammatory cytokines in the anterior, middle and posterior regions of the cortex. n=7 (sham) and 9 (BDL). **(E)**Motor function was assessed by the rotarod performance test. n=9 per group. \**p*<0.05, \*\**p*<0.01, \*\*\**p*<0.001.

## Results

### Basic characteristics

Four weeks after BDL surgery, rats displayed significantly increased spleen/body weight ratio and spleen length compared to sham-operated rats (Supplemental Table 2), suggesting the presence of portal hypertension. In addition, plasma ammonia levels increased in the BDL group. As shown in previous studies^26–28^, BDL rats thus exhibited characteristics of hepatic encephalopathy (HE) related to cirrhosis, including collateral shunting and systemic hyperammonemia.

### Brain inflammation is increased in the cortex in BDL rats

In hepatic encephalopathy, ammonia flows into the brain parenchyma due to systemic hyperammonemia. Ammonia binds to glutamate to form glutamine, a toxic compound, which is thought to contribute to the progression of hepatic encephalopathy^29^ (Figure 1A). We thus quantified ammonia metabolites (Figure 1B), microglia activation (Figure 1C) and pro-inflammatory cytokines (Figure 1D) in the posterior, middle and anterior regions of the cerebral cortex (Supplementary Figure 1).

No significant differences were observed in ammonia levels between the two groups in any of these three regions (Left panel, Figure 1B). In contrast, the level of glutamate, a scavenger of ammonia, was significantly decreased in the anterior and middle regions in BDL rats (nmol/ mg protein, anterior: 348±15 vs. 410±16, *p*=0.02; middle: 341±11 vs. 402±10, *p*=0.001), compared to sham rats with a similar trend in the posterior region as well (320±13 vs. 364±20, *p*=0.073). On the other hand, glutamine levels were significantly increased in the anterior and middle regions in BDL rats (nmol/ mg protein, anterior: 404±23 vs. 286±28, *p*=0.007; middle: 413±29 vs. 300±21, *p*=0.014) with no differences in the posterior region (378±29 vs. 341±19, *p*=0.321).

Microglia activation is an indicator of inflammation and was determined by the endpoint number and branch length of each cell (Figure 1C). Activated cells have fewer endpoints and shorter branch lengths. We observed microglia activation in the middle region of the cortex in BDL rats (endpoint number/cell, 13.0±0.4 vs. 19.2±0.8, *p*<0.001; branch length in μm/cell, 140.4±4.7 vs. 201.9±8.2, *p*<0.001), compared to sham rats with a similar trend in the anterior region (endpoint number/cell, 15.3±0.6 vs. 17.3±0.9, *p*=0.09; branch length in μm/cell, 169.0±6.1 vs. 185.9±10.9, *p*=0.207). Figure 1D shows mRNA expression of pro-inflammatory cytokines in the anterior, middle and posterior regions. IL-1β was significantly increased in all three regions in BDL rats, while INFγ, TNFα and Iba1 were increased only in the middle region. These results demonstrate brain inflammation in hepatic encephalopathy with varying degrees of severity in different regions of the cortex.

### Motor function is impaired in BDL rats

Motor function is controlled by the motor area lying in the anterior and middle regions of the cortex, where we found profound neuroinflammation (Figures 1B - D), and can be evaluated by the rotarod performance test. We found that motor function was significantly reduced in BDL rats (time in seconds, 132±29 vs. 221±18, *p*=0.023), suggesting that increased inflammation in the cerebral cortex contributes to motor function impairment in hepatic encephalopathy (Figure 1E).

### Neuroinflammation occurs in human brains of cirrhotic patients with hepatic encephalopathy

We performed bioinformatic analysis of brains of cirrhotic patients with hepatic encephalopathy using data from GSE41919 (Figure 2A). A quality control analysis identified some samples overlapping between the HE and control groups, resulting in their removal from further analysis (Supplemental Figure 2). Subsequently, hierarchical clustering plots demonstrated that the remaining samples in each group were closely clustered with clear separation between the HE and control groups (Figure 2B). A PCA plot confirmed no overlap between the two groups (Figure 2C). Thus, the HE and control groups analyzed were transcriptionally distinct.

**Figure 2.**
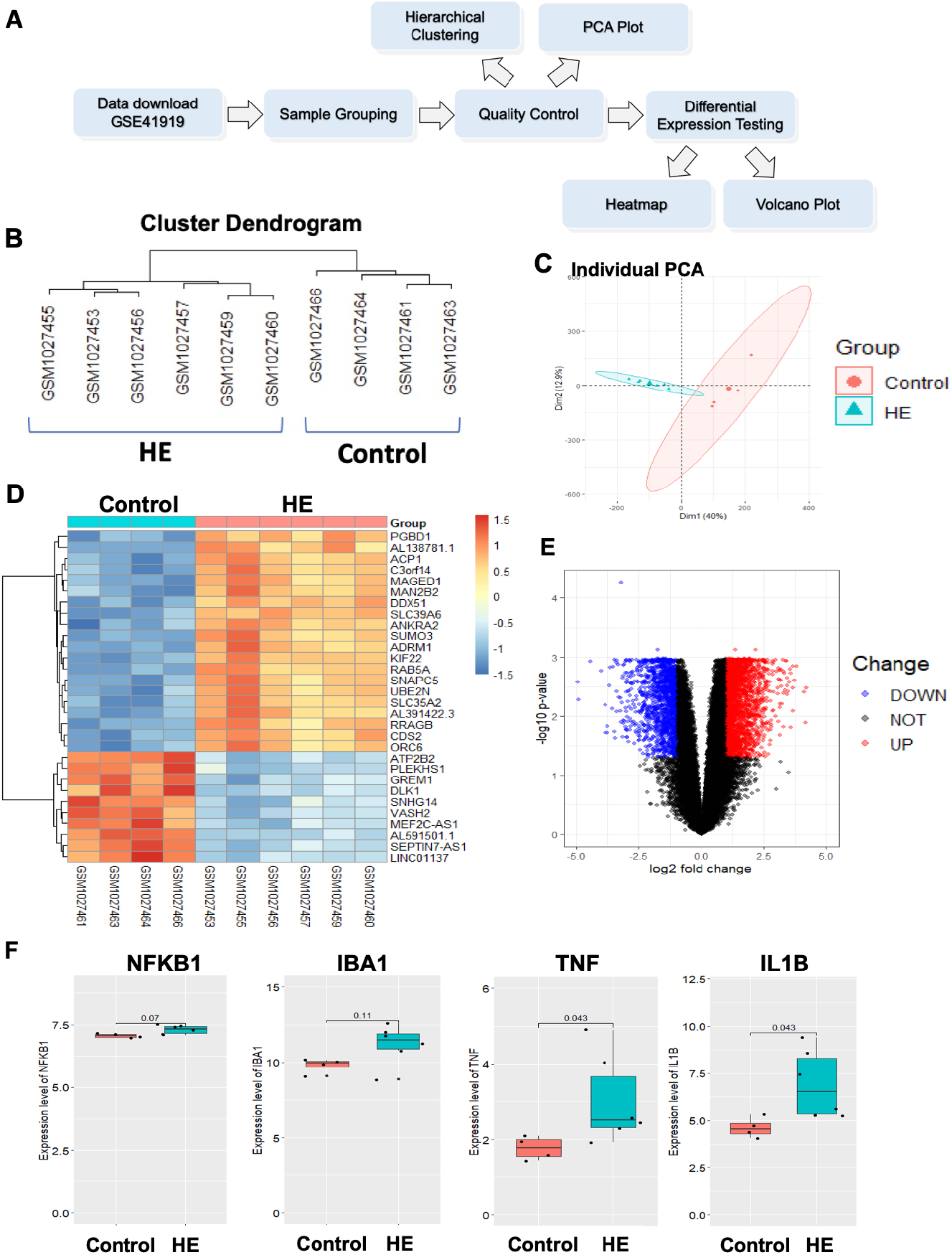
Brain inflammation is also observed in human brains of cirrhotic patients with hepatic encephalopathy. **(A)** A workflow for bioinformatic analysis of human brains of cirrhotic patients with hepatic encephalopathy (HE). **(B)** Hierarchical clustering of samples from the HE and control groups after removing low-quality samples. **(C)** A PCA plot of the GSE41919 dataset after removing low-quality samples. **(D)** A heatmap of the top 30 differentially expressed genes (DEGs) between the HE and control groups in GSE41919. n=4 (control) and 6 (HE). **(E)** A volcano plot of DEGs between the HE and control groups. Red: upregulated genes; blue: downregulated genes. **(F)** Expression levels of NF-κβ, Iba1, TNFα and IL-1β (NFKB1, IBA1, TNF and IL1B in human gene annotation, respectively) in the HE and control groups. n=4 (control) and 6 (HE).

A total of 3800 differentially expressed genes (DEGs) were identified with 2320 genes upregulated and 1480 genes downregulated in the HE group compared to the control group. A heatmap of the top 30 DEGs is presented in Figure 2D. A volcano plot is also provided in Figure 2E to show a comprehensive gene profile, in which red dots represent upregulated genes and blue dots represent downregulated genes. Further, we specifically compared expression levels of several inflammation-related genes between the two groups. We found that NF-κβ, Iba1, TNFα and IL-1β (NFKB1, IBA1, TNF and IL1B in human gene annotation, respectively) were upregulated in the HE group with statistical significance for TNFα and IL-1β (Figure 2F).

### Meningeal lymphangiogenesis is significantly increased in BDL rats

An entire distribution of lymphatic and blood vessels in the meninges is shown in Figure 3A. Whole meninges were stained with Prox1, a transcription factor expressed in lymphatic endothelial cells (white dots), and CD31, an endothelial cell marker (blue). Meningeal lymphatic vessels were found to develop from both lateral sides along with middle meningeal arteries (MMA) and extend toward the edge of the superior sagittal sinus (SSS) and transverse sinus (TS) without attaching to blood vessels. Meningeal lymphangiogenesis was defined as meningeal lymphatic vascular growth expressed by the ratio of the lymphatic vascular length to the meningeal area surrounded by the cerebrum olfactory bulb fissure (COF), SSS and TS. BDL rats exhibited significantly increased lymphangiogenesis compared to sham rats (0.0017±0.0001 vs. 0.0011±0.0002, *p*=0.02) (Figure 3B).

**Figure 3.**
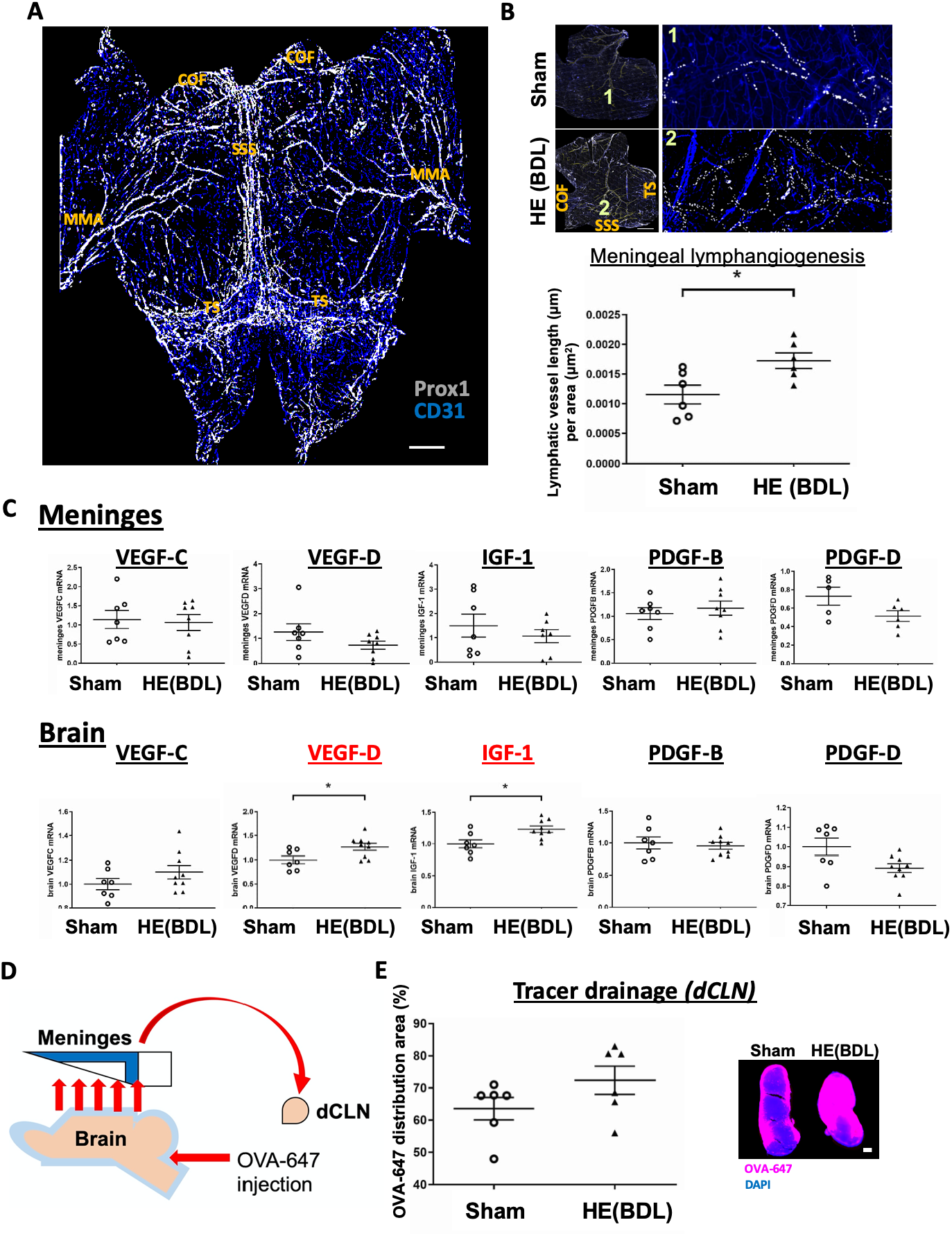
Lymphangiogenesis in the meninges is significantly increased in BDL rats. **(A)** Rat meninges were dissected from the skull. The cerebrum olfactory bulb fissure (COF), transverse sinus (TS) and middle meningeal artery (MMA) are located in the anterior, posterior and lateral edges of the meninges, respectively. Lymphatic endothelial cells were immunolabeled with Prox1 (white), while CD31 (blue) was used as an endothelial cell marker. Scale bar: 2mm. **(B)** Lymphatic vessels were depicted by connecting Prox1 positive nuclei. Lymphatic vessels in the area surrounded by the COF, SSS and TS were analyzed. Lymphangiogenesis was defined by the ratio of the lymphatic vessel length to the meningeal area analyzed. Scale bar: 2mm. n=6 per group. \**p*<0.05. **(C)** mRNA expression of lymphangiogenic factors in the meninges and brain. n=7 (sham) and 9 (BDL). \**p*<0.05. **(D)** A schematic of OVA-647 injection. OVA-647 injected into the cisterna magna leaves the brain parenchyma and returns to the sub-arachnoid space. It then flows to the meninges and drains to the deep cervical lymph nodes (dCLN) through meningeal lymphatic vessels. **(E)** OVA-647 drainage into the dCLN of sham and BDL rats as an indicator of meningeal lymphatic drainage. Scale bar: 200μm.

We then examined expression levels of lymphangiogenic factors in the meninges and brain (Figure 3C). While none of the factors differed between the two groups in the meninges, the brain showed significantly increased expression of VEGF-D (1.26±0.07 vs. 1.00±0.08, *p*=0.028) and insulin like growth factor-1 (IGF-1) (1.23±0.05 vs. 1.00±0.06, *p*=0.012) in BDL rats compared to sham rats with a similar trend for VEGF-C. These observations may indicate that lymphangiogenic factors are generated in the brain and released into the meninges to induce lymphangiogenesis.

To assess meningeal lymphatic drainage in BDL rats, we injected a tracer dye (OVA-647) into the cisterna magna and examined the OVA-647 distribution in the deep cervical lymph nodes (dCLN) (Figure 3D). OVA-647 in the subarachnoid space flows to the meninges and then to the dCLN through meningeal lymphatic vessels. Some OVA-647 was taken up by macrophages in the meninges before reaching meningeal lymphatic vessels (Supplemental Figure 3). OVA-647 drainage to the dCLN tended to increase in BDL rats (%, 72.4±4.4 vs. 63.6±3.5, *p*=0.146) (Figure E).

### VEGF-C-induced meningeal lymphangiogenesis increases meningeal lymphatic drainage in BDL rats

Adeno-associated virus (AAV)8-VEGF-C or AAV8-GFP (control) was injected into the cisterna magna at a dose of 1.5 x 10^11^ GC/rat one day after BDL surgery to locally express VEGF-C and thus to increase meningeal lymphangiogenesis (Figure 4A). Four weeks after BDL surgery (3 weeks and 6 days after AAV injection), there was a significant 10-fold increase in VEGF-C mRNA expression in the meninges of rats injected with AAV8-VEGF-C (28.8±9.1 vs. 2.9±1.0, *p*=0.022), compared to those injected with control AAV8. As indicated by GFP, strong AAV8-derived gene expression was observed in the COF and TS areas in the meninges while the brain showed weak expression in the cerebellum and even weaker expression in the cortex (Supplemental Figure 4). Consistent with increased VEGF-C mRNA expression, meningeal lymphangiogenesis was increased 1.5-fold in BDL rats with AAV8-VEGF-C (*p*=0.035) (Figure 4B). There were no differences in body weight, spleen/body weight ratio, spleen length or plasma ammonia levels between control and VEGF-C treated BDL rats (Supplemental Table 3).

**Figure 4.**
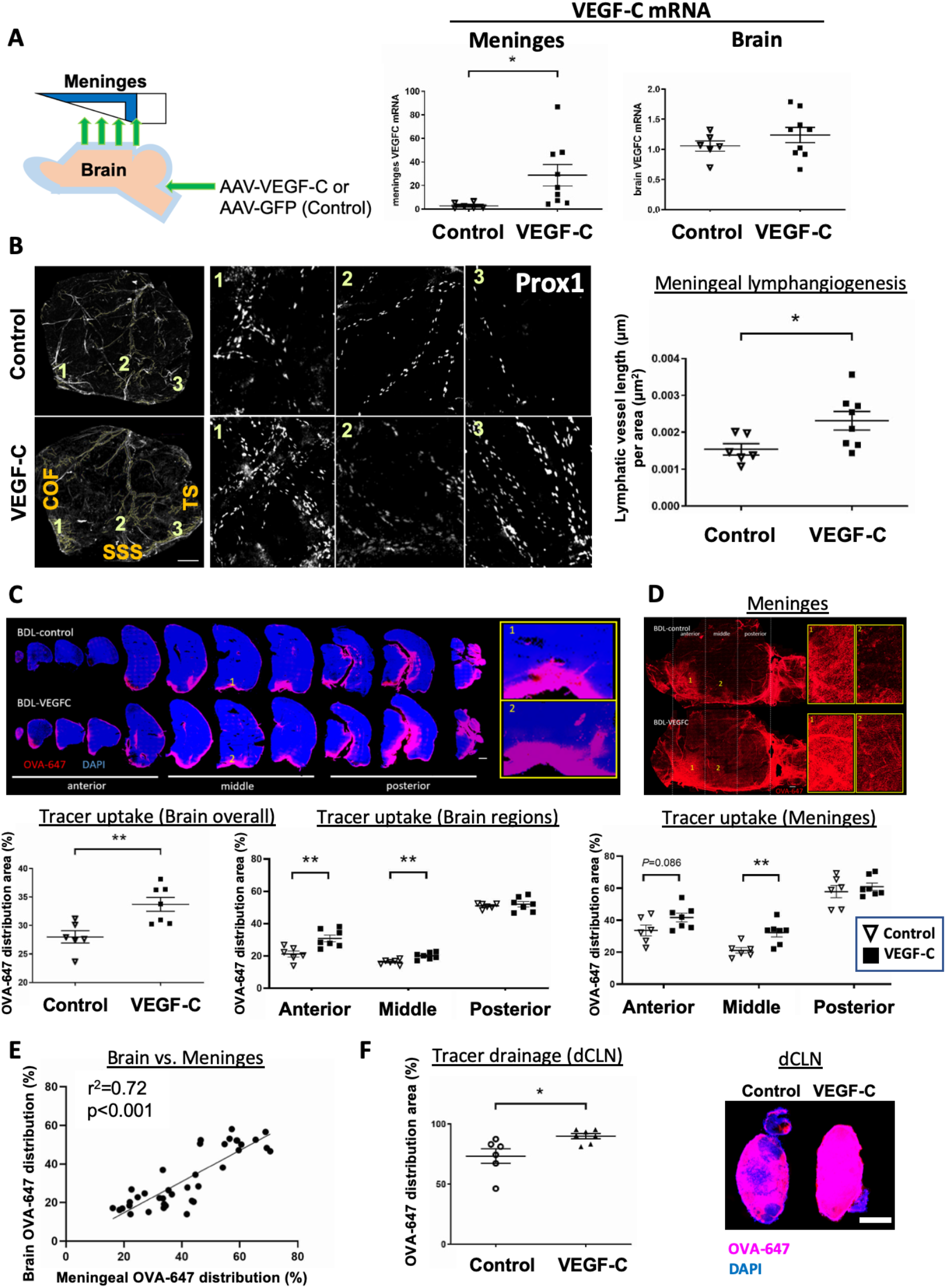
VEGF-C increase meningeal lymphangiogenesis and improves brain glymphatic function and meningeal lymphatic drainage in BDL rats. **(A)** A schematic of AAV8 injection and VEGF-C mRNA expression in the meninges and brain. AAV8-VEGF-C or AAV8-GFP (control) was injected into the cisterna magna at a dose of 1.5×10^11^ GC/rat one day after BDL surgery. Four weeks after BDL surgery, samples were collected for analysis. n=6 (control) and 9 (VEGF-C). **(B)** Meningeal lymphagiogenesis. White dots: Prox1 (a marker of lymphatic endothelial cells). Vessel length was determined as an indicator of lymphangiogenesis. COF: cerebrum olfactory bulb fissure; SSS: superior sagittal sinus; TS: transverse sinus. Scale bar: 2mm. n=6 (control) and 9 (VEGF-C). **(C)** Brain glymphatic function. OVA-647 was injected into the cisterna magna and its positive areas were determined as an indicator of brain glymphatic function. Scale bar: 1mm. n=6 (control) and 9 (VEGF-C). **(D)** Meningeal lymphatic function. OVA-647 positive areas were used as an indicator of meningeal lymphatic function. Scale bar: 1mm. n=6 (control) and 9 (VEGF-C). **(E)** Brain glymphatic function was positively correlated with meningeal lymphatic function. **(F)** Drainage to the deep cervical lymph nodes (dCLN). OVA-647 intensity was used as an indicator of meningeal lymphatic drainage. Scale bar: 1mm. \**p*<0.05, \*\**p*<0.01.

We also determined effects of VEGF-C overexpression on tracer uptake in the brain and meninges as well as meningeal lymphatic drainage in BDL rats (Figures 4C - F). We found significantly increased tracer uptake in the brain in the VEGF-C group (%, 33.7±1.2 vs. 28.0±1.1, *p*=0.005), particularly in the anterior (30.9±2.1 vs. 21.2±1.9, *p*=0.006) and middle regions (20.1±0.8 vs. 16.3±0.6, *p*=0.003) of the cortex, compared to the control group (Figure C). There were no differences in the posterior region (*p*=0.606). Similar to the brain, VEGF-C overexpression enhanced meningeal tracer uptake in the anterior (%, 41.7±2.7 vs. 33.6±3.4, *p*=0.086) and middle regions (%, 32.2±2.7 vs. 21.1±1.7, *p*=0.006) (Figure 4D), compared to the control group, with no differences in the posterior region. Further, brain glymphatic function was positively correlated with meningeal lymphatic function (r^2^=0.720, *p*<0.001) (Figure 4E). Furthermore, OVA-647 drainage in the deep cervical lymph nodes (dCLN) was significantly increased in the VEGF-C group (%, 89.9±2.1 vs. 73.4±6.1, *p*=0.02) (Figure 4F). Collectively, these findings strongly suggest that VEGF-C overexpression, through increased meningeal lymphangiogenesis, facilitates meningeal lymphatic drainage in hepatic encephalopathy.

### VEGF-C-induced meningeal lymphangiogenesis attenuates brain inflammation and restores motor function in BDL rats

VEGF-C did not change ammonia, glutamate or glutamine levels in the cerebral cortex in BDL rats (Figure 5A). However, VEGF-C attenuated microglia activation in the anterior and middle regions of the cortex as indicated by increased endpoint numbers/cell in the anterior (21.8±0.9 vs. 15.8±0.6, *p*<0.001) and middle regions (22.8±0.6 vs. 15.8±1.0, *p*<0.001) as well as increased branch lengths (μm/cell) in the anterior (238.9±9.3 vs. 162.9±4.2, *p*<0.001) and middle regions (243.0±8.0 vs. 165.4±10.0, *p*<0.001) (Figure 5B). Microglia activation did not differ in the posterior region between the two groups.

**Figure 5.**
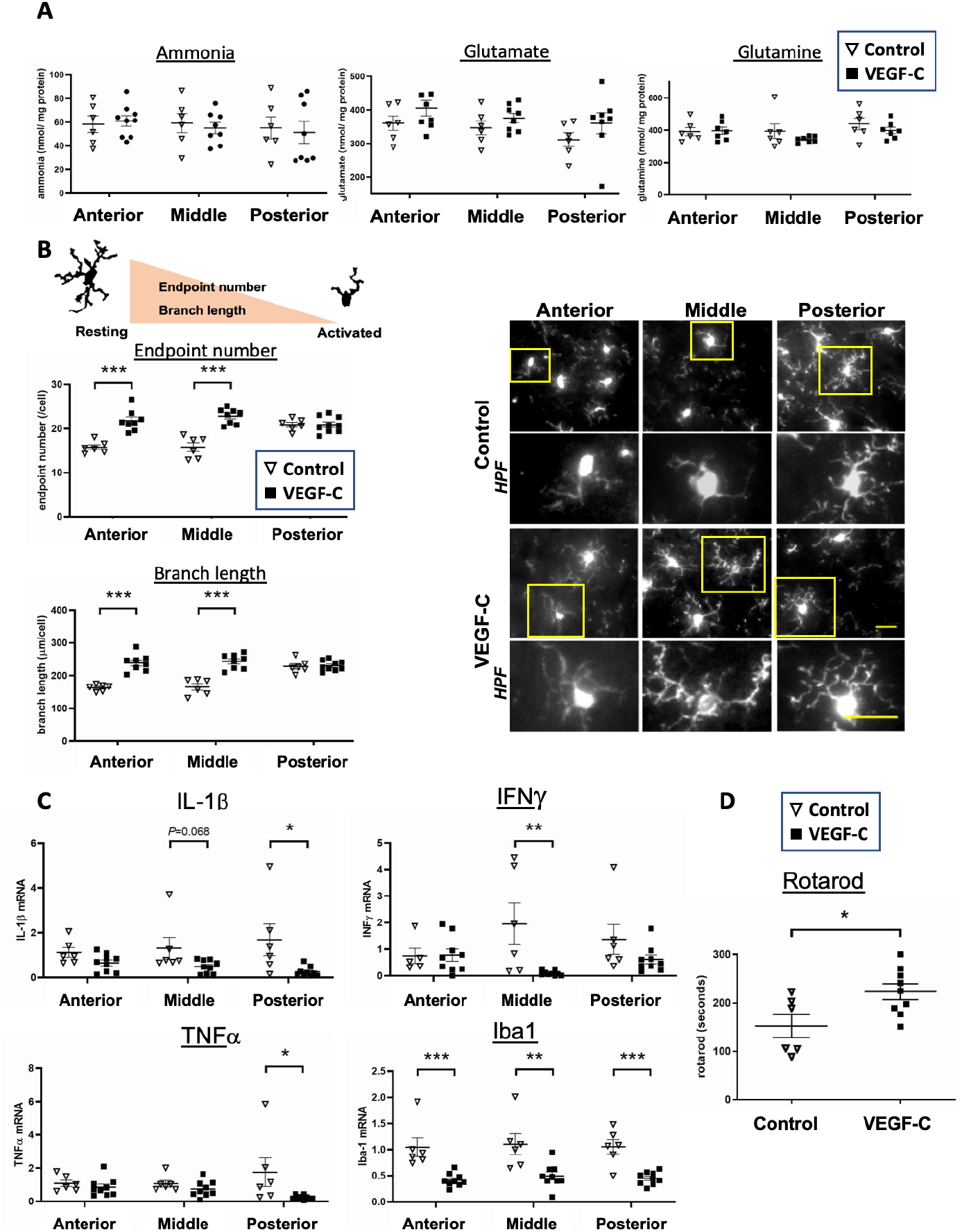
VEGF-C attenuates brain inflammation and improves motor function in BDL rats. AAV8-VEGF-C or AAV8-GFP (control) was injected into the cisterna magna at a dose of 1.5×10^11^ GC/rat one day after BDL surgery. Four weeks after BDL surgery, sample collection was performed. The rotarod performance test was conducted one day before sample collection. n=6 (control) and 9 (VEGF-C) for all experiments. **(A)** Ammonia metabolites in the anterior, middle and posterior regions of the cortex. **(B)** Microglia activation determined by the endpoint number and branch length of each cell. Microglia were immunolabeled with Iba1. Scale bar: 25μm. **(C)** mRNA expression of pro-inflammatory genes in the three regions of the cortex. **(D)** Motor function assessed by the rotarod performance test. \**p*<0.05, \*\**p*<0.01, \*\*\**p*<0.001.

VEGF-C also attenuated expression of pro-inflammatory genes in the anterior, middle or posterior region of the cortex in BDL rats (Figure 5C). IL-1β tended to decrease in all regions of the cortex in the VEGF-C group (anterior: 0.65±0.13 vs. 1.12±0.22, *p*=0.075; middle: 0.49±0.10 vs. 1.31±0.49, *p*=0.068; and posterior: 0.26±0.07 vs. 1.69±0.71, *p*=0.027), compared to the control group. Expression levels of INFγ and TNFα were significantly decreased in the middle and posterior regions in VEGF-C treated BDL rats, respectively. Iba1 expression was significantly reduced in all regions (anterior: 0.42±0.04 vs. 1.05±0.18, *p*=0.001; middle: 0.50±0.08 vs. 1.10±0.20, *p*=0.006; and posterior: 0.45±0.04 vs. 1.05±0.14, *p*<0.001). We also demonstrated that VEGF-C overexpression significantly improved motor function in BDL rats (time in seconds, 223±16 vs. 152±24, *p*=0.024), compared to those treated with control AAV (Figure 5D). Collectively, these results strongly suggest that VEGF-C overexpression in the brain ameliorates brain inflammation by facilitating meningeal lymphatic drainage and improves hepatic encephalopathy.

### Identification of genes and pathways in the middle cortex restored by VEGF-C-induced meningeal lymphangiogenesis

RNA-seq analysis identified 283 differentially expressed genes (DEGs) between the BDL and sham groups with 170 genes upregulated and 113 genes downregulated in the BDL group. Further, a group of BDL rats treated with AAV8-VEGF-C showed 69 upregulated and 99 downregulated genes, compared to a group of BDL rats with control AAV. Heatmaps of the top 30 DEGs in the comparisons of sham vs. BDL and control vs. VEGF-C are presented in Supplemental Figure 5. A Venn diagram showed an overlap of 21 genes (Figure 6A) and their gene expression profiles are presented in Figure 6B. Gene Set Enrichment Analysis (GSEA) revealed that VEGF-C treatment could activate the proteasome pathway and suppress NF-κβ signaling, Fc gamma R-mediated phagocytosis, phagosome, antigen processing and presentation, cell adhesin molecules, endocytosis, and glutamatergic synapse (Figure 6C). Decreased NF-κβ signaling is consistent with attenuated brain inflammation observed in VEGF-C treated BDL rats. Decreased phagocytosis and phagosome may also be related to decreased microglia activation by VEGF-C, reflecting improved brain inflammation.

**Figure 6.**
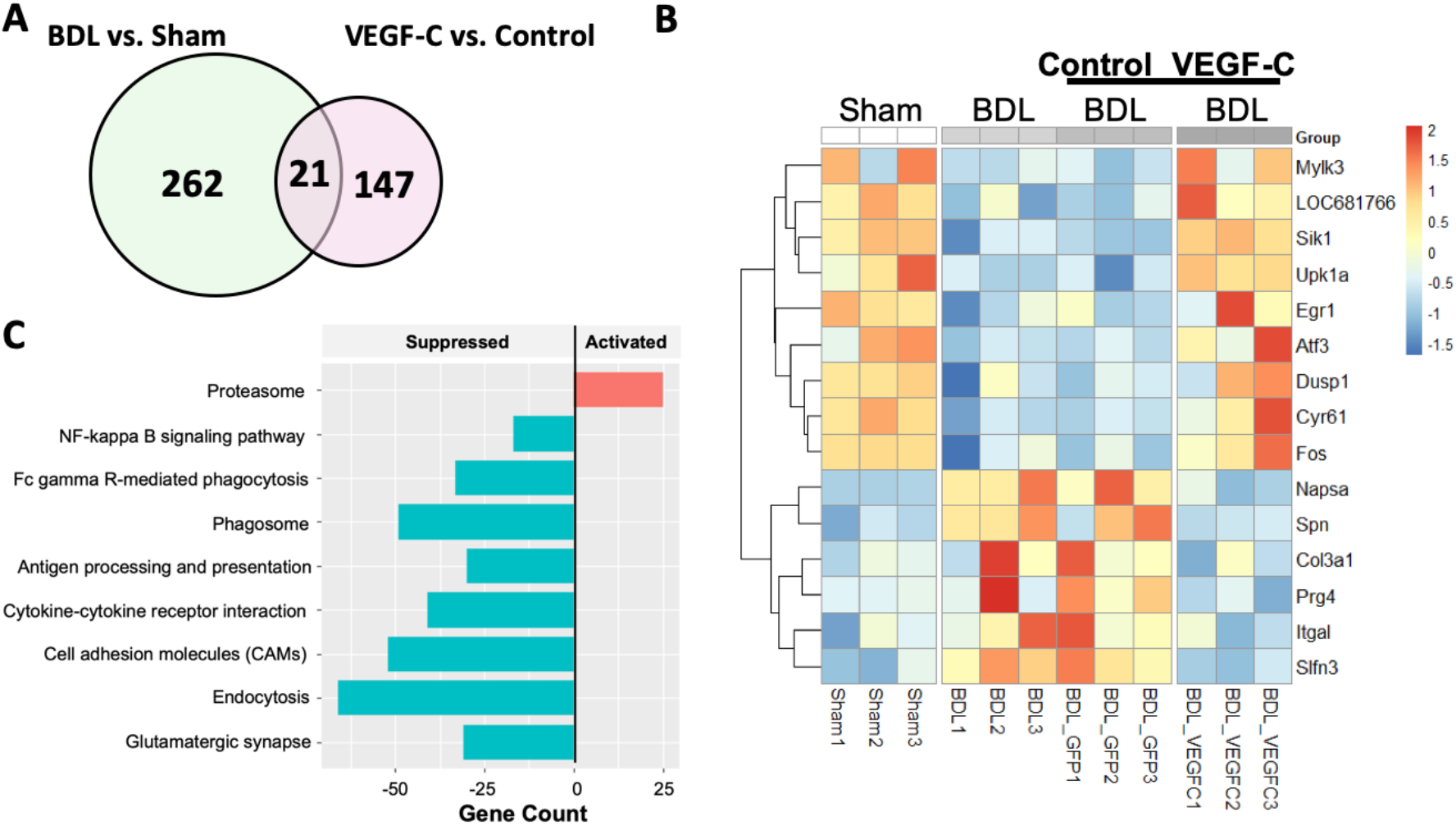
The effect of VEGF-C overexpression on the gene expression profile and functional changes in the middle region of the cortex. RNA-seq analysis. n=3 per group. **(A)** A Venn diagram reveals intersected genes. **(B)** A heatmap shows differentially expressed genes among the groups to identify genes that were altered by BDL but recovered to the levels of the sham group by VEGF-C overexpression. **(C)** Gene Set Enrichment Analysis (GSEA) was conducted to examine functional changes caused by VEGF-C overexpression.

## Discussion

Our study has demonstrated the importance of the meningeal lymphatic system for the treatment of HE. We showed that VEGF-C overexpression increases meningeal lymphangiogenesis and enhances meningeal lymphatic drainage, leading to decreased brain neuroinflammation and improved motor function in HE. To our knowledge, this is the first demonstration that increased meningeal lymphatic drainage could decrease neuroinflammation (e.g., microglial activation and pro-inflammatory cytokine expression). While most studies of HE focus on its therapeutic targets outside of the central nervous system, such as modulating ammonia levels in the circulation and modifying the gut microbiome, our study presents a new modality of HE therapy with enhanced meningeal lymphatic drainage.

Hyperammonemia has been appreciated for years as a key pathogenic mechanism of HE. Infection and resulting systemic inflammation are also the underlying precipitants of many episodes of HE, and a proinflammatory state in cirrhotic patients has been shown to work synergistically with hyperammonemia to impair neurologic function^30^. However, targeting solely these events may not be sufficient to reduce the accumulation of harmful substances in the brain and cure inflammation and HE. This is because the pathogenesis of HE is multifactorial, as reflected in observations that ammonia levels are elevated in many cirrhotic patients without HE and often do not normalize with resolution of HE^31–33^. In the current study, VEGF-C-induced increased meningeal lymphangiogenesis and drainage ameliorated brain inflammation and motor dysfunction in HE. Some very recent studies have also reported the benefits of manipulation of meningeal lymphatic drainage^12, 34, 35^. In these studies, increased meningeal lymphatic drainage resulted in a better response to immune checkpoint therapy and eradicating brain tumors^34, 35^, as well as contributing to improved learning and memory performance in age-associated neurological disorders such as Alzheimer’s disease^12^. Thus, a combination of lowering plasma ammonia and facilitating meningeal lymphatic drainage could be a more effective strategy for the treatment of HE.

We investigated the potential mechanism of how increased meningeal lymphatic drainage ameliorates neuroinflammation in HE using an unbiased transcriptomic approach with a focus on the middle region of the cortex in all groups examined in this study. The middle region of the cortex was chosen because microglial activation and increased expression of pro-inflammatory cytokines induced by BDL were most prominent in this region among the three regions analyzed. We identified genes significantly altered in HE, but restored by VEGF-C treatment. Among them, salt-inducible kinase 1 (Sik1), expressed in microglia, may be noteworthy. Sik1 is a serine/threonine protein kinase^36^ and its upregulation was shown in the rat cortex and hippocampus in response to kainic acid-induced seizures^37^ as well as in the rat brain in response to alcohol intake^38^. In the latter study^38^, knockdown of Sik1 in cultured microglia activated the NFkB pathway and induced apoptosis, suggesting that Sik1 protects microglia from inflammation and apoptosis. In our bioinformatic analysis, a significant decrease in Sik1 expression was observed in BDL rats, but was restored by VEGF-C overexpression to a level similar to that of sham rats. Further, pathway analysis indicated that VEGF-C treatment significantly suppressed NFkB signaling in the middle region of the cortex in BDL rats. These observations may suggest that VEGF-C-induced increased lymphatic drainage restored Sik1 expression, leading to inhibition of the NFkB pathway in the brain, and that modulation of Sik1 could be a potential therapeutic option for the treatment of neuroinflammation. Further investigations are necessary to test this possibility. Nevertheless, our study demonstrated for the first time that increased meningeal drainage could reduce neuroinflammation (e.g., microglia activation) and HE.

Although the occurrence of neuroinflammation in HE is known, it is poorly understood which inflammatory cytokines are specifically upregulated. We have specified significant upregulation of TNFα, IL-1β and Iba1 in BDL rats. Importantly, our bioinformatic analysis of human brains also revealed significant increases of TNFα and IL-1β in cirrhotic patients with HE as well as a trend toward increase in Iba1. To our knowledge, this may be the first report to show upregulation of specific inflammatory genes in humans in association with HE. In addition, we have demonstrated that neuroinflammation and microglia activation occurred to different degrees in different regions of the cerebral cortex, with the most significant degree in the middle region. Given a great degree of division of functional specificity of the brain, the middle region may be particularly related to the pathogenesis of HE. The detailed regional data of neuroinflammation this study provides are valuable for the study of HE and its treatment.

Lymphangiogenesis is increased in many pathological conditions in many organs^18, 39^. We have also observed significantly increased lymphangiogenesis in the meninges of cirrhotic rats with HE (BDL rats). While the underlying mechanism requires further investigation, our results suggest that lymphangiogenic factors generated in the brain, such as VEGF-D and IGF-1, may be transported to the meninges and induced lymphangiogenesis. This is based on our observation that none of well-known lymphangiogenic genes were upregulated in the meninges of BDL rats, while VEGF-D and IGF-1 showed significant increases in their expression in the brain. VEGF-C exhibited a trend toward upregulation, but its increase was not statistically significant. Interestingly, intra-cisternal injection of AAV-VEGF-C and control AAV-GFP to BDL rats resulted in larger increases in expression of these genes in the meninges than the brain 4 weeks after their injection, while acute injection of a tracer dye OVA-647 to BDL rats showed its uptake by both the brain and the meninges. These observations may suggest transport of substances from the brain to the meninges over a period of time, including injected VEGF-C and GFP genes and/or their protein products.

Brain edema, characterized by an excessive accumulation of fluid in the intracellular or extracellular space of the brain, is commonly observed in patients with HE^40^. Generally in peripheral tissues, edema is a strong inducer of lymphangiogenesis to clear excessive interstitial fluid. A relationship between cerebral edema and the development of meningeal lymphangiogenesis would be an interesting area of investigation, as it may lead to important discoveries about the mechanism of meningeal lymphangiogenesis and its role in the pathogenesis of HE.

Despite increased meningeal lymphangiogenesis in BDL rats with HE, no significant differences were observed in drainage to the deep cervical lymph nodes (dCLN) between the BDL and sham groups (although a trend of increased drainage was found in the BDL group). It is possible that these meningeal lymphatic vessels may not be fully functional due to the disease condition of BDL rats. It is reported that lymphatic vessel function is impaired in aging mice ^11, 12^. It is also known that phenotypic changes of lymphatic endothelial cells often occur in diseased tissues^41, 42^. Insufficient meningeal lymphatic drainage function may mediate increased brain inflammation in rats with HE. Examining a possibility of dysfunctional drainage in HE will also be important to understanding the role of the meningeal lymphatic system and the functional importance of meningeal lymphangiogenesis in HE.

In conclusion, the meningeal lymphatic system may become incapable of clearing increased harmful metabolites in the brain in cirrhosis, thereby contributing to the pathogenesis of HE. However, by promoting meningeal lymphangiogenesis via VEGF-C overexpression and enhancing waste clearance, HE could be ameliorated. Manipulation of meningeal lymphangiogenesis could be a new therapeutic strategy for the treatment of HE.

## Abbreviations

HE: hepatic encephalopathy
BDL: bile duct ligation
AAV: adeno-associated virus
VEGF-C: vascular endothelial growth factor-C
IL-1β: interleukin-1beta
INFγ: interferon gamma
TNFα: tumor necrosis factor alpha
CNS: central nervous system
CSF: cerebral spinal fluid
dCLN: deep cervical lymph node
BBB: brood-brain barrier
VEGFR3: vascular endothelial growth factor receptor 3
COF: cerebrum olfactory bulb fissure
SSS: superior sagittal sinus
TS: transverse sinus
TMM: trimmed mean of means
GSEA: Gene Set Enrichment Analysis
FDR: false discovery rate
PCA: principal component analysis
DEG: differentially expressed gene
FC: fold change
SEM: standard error of the mean
NFkB: nuclear factor kappa-light-chain-enhancer of activated B cell
Iba1: ionized calcium binding adaptor molecule 1
MMA: middle meningeal artery
Sik1: salt-inducible kinase 1
IGF-1: insulin-like growth factor 1

## Acknowledgements

We would like to thank Dr. Jeffery Kocsis (Yale University) for technical support, and Drs. Anne Eichmann, Jean-Leon Thomas, and Helene Benveniste (Yale University) for valuable discussions.

