## Supplemental Materials for "Enhanced meningeal lymphatic drainage ameliorates neuroinflammation and hepatic encephalopathy in cirrhotic rats"

**Content:**

1. **Supplemental Figures: 5**
2. **Supplemental Tables: 3**
3. **Supplemental Methods**

**Supplemental Figures**

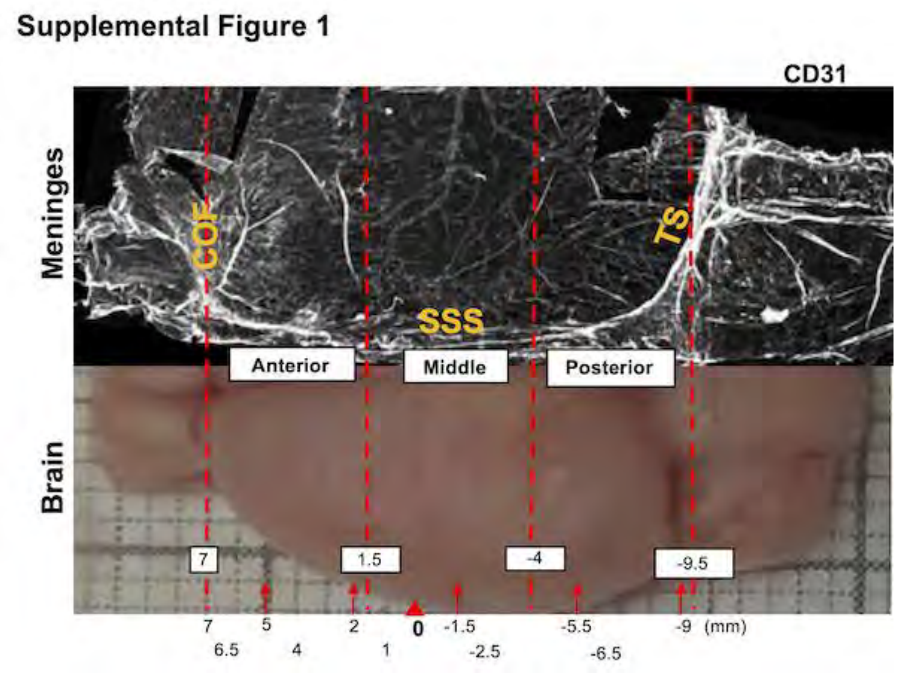


**Supplemental Figure 1. Definition of brain sections and the anterior, middle and posterior regions of the brain and meninges in the rat.**

To evaluate brain glymphatic function and meningeal lymphatic function, we examined the distribution of OVA-647 tracer in 11 brain slices positioned at 7, 6.5, 5, 4, 2, 1, -1.5, -2.5, -5.5, -6.5 and -9 mm relative to the bregma. We also defined the anterior, middle and posterior regions in the brain and meninges based on their position at 7, 1.5, -4 and -9.5 mm relative to the bregma. The width of each region was identical (5.5 mm). Grid: 1mm. COF: cerebrum olfactory bulb fissure; TS: transverse sinus; SSS: superior sagittal sinus. White lines shown in the meninges are blood and lymphatic vessels immunolabeled with CD31 antibody.

### Supplemental Figure 2

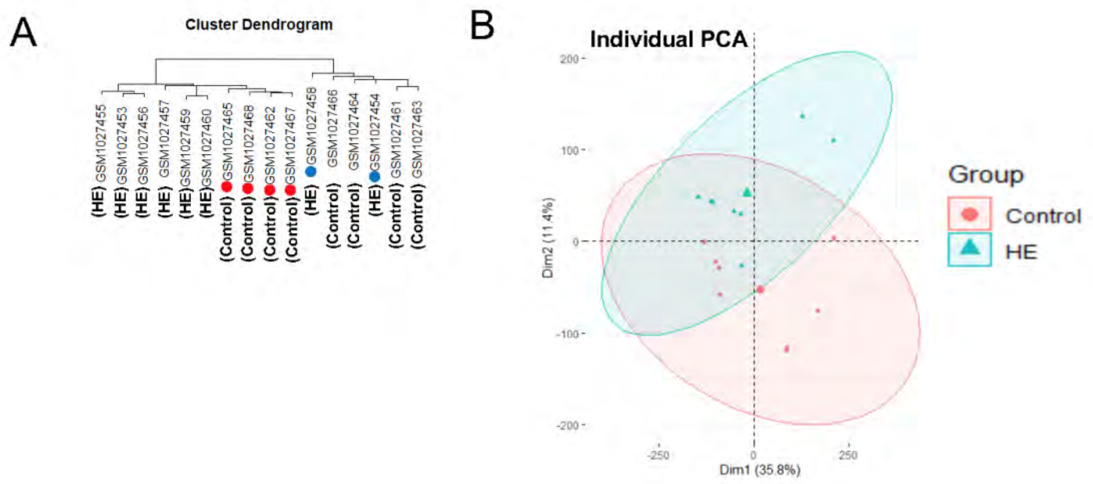


**Supplemental Figure 2. A quality control analysis for sample selection.**
**(A)** Hierarchical clustering of samples from hepatic encephalopathy (HE) and control groups before removing low-quality samples. Four samples from the control group (marked in red) were clustered to the HE group, while 2 samples from the HE group (marked in blue) were clustered to the control group, indicating a quality issue of these samples and resulting in their removal from our analysis. **(B)** A PCA plot of the GSE41919 dataset before removing low-quality samples. The circles originated from the HE and control groups were overlapped in the PCA plot, confirming the above finding.

Supplemental Figure 3

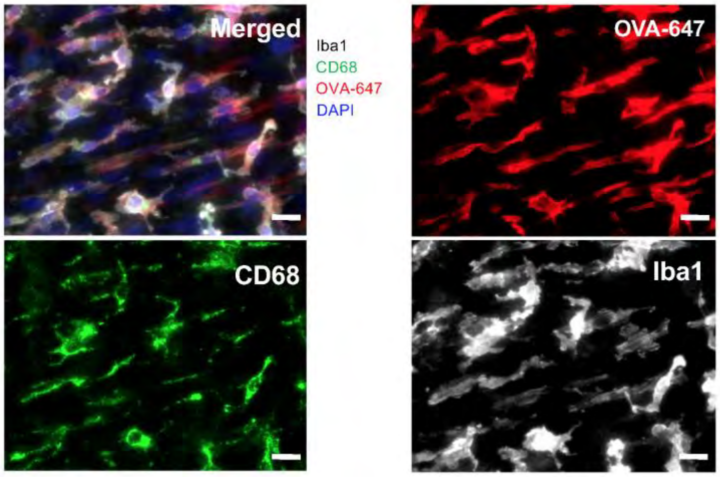


**Supplemental Figure 3. OVA-647 tracer uptake by macrophages in the meninges.** Macrophages immunolabeled with Iba1 and CD68 were colocalized with a tracer dye OVA-647, indicating that macrophages take up OVA-647 in the meninges. Scale bar: 20μm.

### Supplemental Figure 4

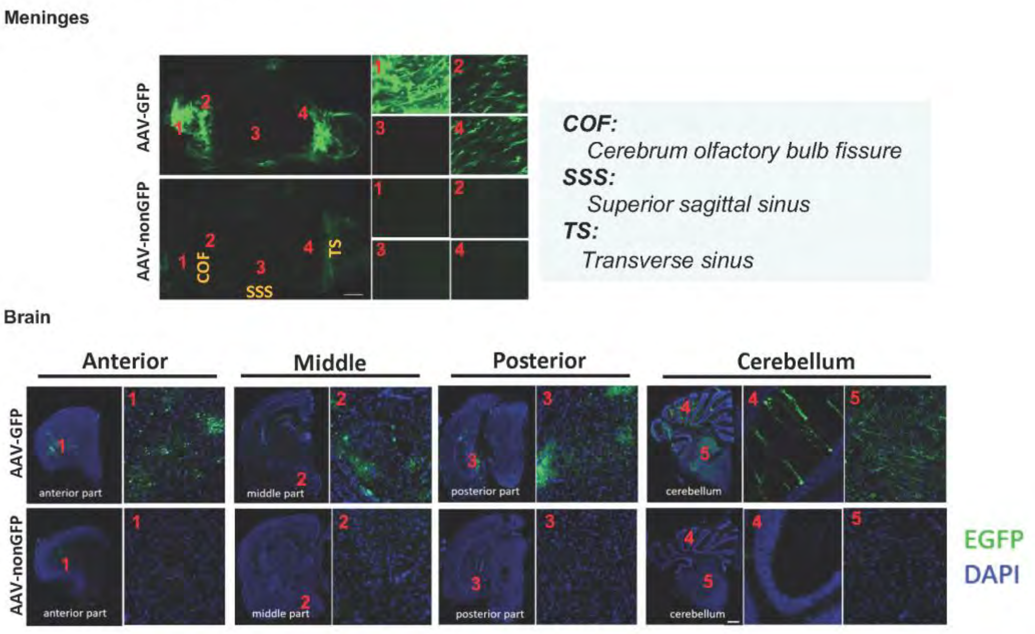


**Supplemental Figure 4. GFP expression in the meninges and brain 4 weeks after BDL.** AAV8-GFP was injected into the cisterna magna at a dose of $1.5x10^{11}$ GC/rat one day after BDL surgery. Four weeks after BDL surgery, strong GFP expression was found in the COF and TS areas of the meninges with weaker expression in the cortex and cerebellum areas of the brain, indicating successful AAV infection. COF: cerebrum olfactory bulb fissure; TS: transverse sinus; SSS: superior sagittal sinus. Scale bars: meninges, 1mm; brain, 2mm.

### Supplemental Figure 5

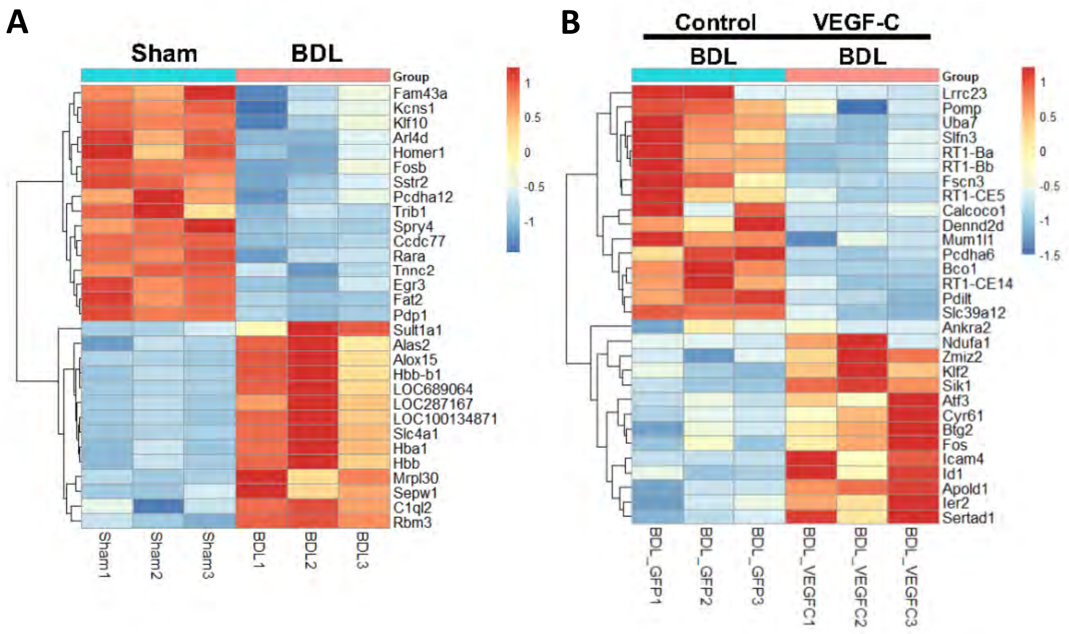


**Supplemental Figure 5. Heatmaps of the top 30 differentially expressed genes.**

**(A)** Sham vs. BDL group. **(B)** BDL: GFP-AAV (control) vs. BDL: VEGF-C-AAV treated group.

**Supplemental Table 1. Primer sequences**

| *Gene name* | *GeneBank accession no.* | | *Primer sequences (5'-3')* | *Expected size (bp)* | *Annealing temperature for qPCR (°C)* |
|---|---|---|---|---|---|
| **Iba1** | **NM_017196.3** | ***F:*** | **CAAGGATTTGCAGGGAGGAA** | **97** | **60** |
| | | ***R:*** | **CTTGGGATCATCGAGGAAGTG** | | |
| **IL1β** | **NM_031512.2** | ***F:*** | **CTATGGCAACTGTCCCTGAA** | **113** | **60** |
| | | ***R:*** | **GGCTTGGAAGCAATCCTTAATC** | | |
| **IFNγ** | **NM_138880.2** | ***F:*** | **GTTTCCCAAGGACGGTAACA** | **96** | **60** |
| | | ***R:*** | **CTGATGGCCTGGTTGTCTTT** | | |
| **TNFα** | **NM_012675.3** | ***F:*** | **GCCTCAGCCTCTTCTCATTC** | **100** | **60** |
| | | ***R:*** | **GGGAACTTCTCCTCCTTGTTG** | | |
| **GFAP** | **NM_017009.2** | ***F:*** | **TGGTATCGGTCCAAGTTTGC** | **99** | **60** |
| | | ***R:*** | **TTGGCGGCGATAGTCATTAG** | | |
| **MCP** | **NM_031530.1** | ***F:*** | **GTCTCAGCCAGATGCAGTTAAT** | **105** | **60** |
| | | ***R:*** | **CTGCTGGTGATTCTCTTGTAGTT** | | |
| **VEGFC (rat)** | **NM_053653.1** | ***F:*** | **TGCCGGTGCATGTCTAAA** | **90** | **60** |
| | | ***R:*** | **CTGCCTGACACTGTGGTAAT** | | |
| **VEGFD** | **NM_031761.1** | ***F:*** | **TGGGACAGAAGACCACTCTTA** | **106** | **60** |
| | | ***R:*** | **TCCAGGACATGGTGCTTTAC** | | |
| **IGF1** | **NM_001082477.2** | ***F:*** | **TCTGCTTGCTCACCTTTACC** | **124** | **60** |
| | | ***R:*** | **CCTGTGGGCTTGTTGAAGTA** | | |
| **FGF2** | **NM_019305.2** | ***F:*** | **GAACCGGTACCTGGCTATGA** | **182** | **60** |
| | | ***R:*** | **CCGTTTTGGATCCGAGTTTA** | | |
| **PDGFD** | **NM_023962.2** | ***F:*** | **CGGCTCATCTTAGTCTCCATTC** | **126** | **60** |
| | | ***R:*** | **GTGATTGCTCTCATCTCTCCTG** | | |
| **PDGFB** | **NM_031524.1** | ***F:*** | **AAAGGGAAGCACCGAAAGT** | **102** | **60** |
| | | ***R:*** | **TAAATAACCCTGCCCACACTC** | | |
| **VEGFC (mouse)** | **NM_009506.2** | ***F:*** | **TGTCTCTGGCGTGTTCCCT** | **190** | **60** |
| | | ***R:*** | **ATCAGCTCATCTACGCTGGAC** | | |
| **18S** | **NR_003278** | ***F:*** | **ACGGAAGGGCACCACCAGGA** | **127** | **60** |

**Supplemental Table 2. Baseline characteristics of sham and cirrhotic rats**

| | Sham | BDL |
|---|---|---|
| **Total numbers$** | n=20 | n=23 |
| **Body weight (g)#** | 369±9 | 355±11 |
| **Spleen/body weight (%)#** | 0.20±0.01 | 0.65±0.03** |
| **Spleen length (cm)#** | 3.7±0.1 | 5.6±0.2** |
| **Plasma ammonia (μM)‡** | 110±26 | 379±67* |

BDL: bile duct ligation.
$: The total number of rats used for analysis. Body, liver and spleen weights as well as plasma ammonia were not measured for all rats used for analysis.
#: n=13 (sham) and 12 (BDL).
‡: n = 6 (sham) and 9 (BDL).
*$p$<0.01, **$p$<0.001.

**Supplemental Table 3. Baseline characteristics of cirrhotic rats treated with AAV8-GFP (Control) or AAV8-VEGF-C**

| | BDL (Control) | BDL (VEGF-C) |
|---|---|---|
| **Total number** | n=6 | n=9 |
| **Body weight (g)** | 362±9 | 352±9 |
| **Spleen/body weight (%)** | 0.72±0.04 | 0.68±0.03 |
| **Spleen length (cm)** | 5.4±0.2 | 5.5±0.1 |
| **Plasma ammonia (μM)** | 307±53 | 324±48 |

BDL: bile duct ligation.

### Supplemental Methods

***Tissue preparation for evaluation of brain glymphatic function and meningeal lymphatic drainage***

Two separate sets of BDL and sham rats were prepared for isolation of the brain and meninges with one set for histological analysis and the other for biochemical analysis. For BDL rats given AAV8-VEGF-C or AAV8-GFP, the brain and meninges isolated were cut into the left and right parts from the midline with the left half for histological analysis and the right half for biochemical analysis.

For histological analysis, the brain was fixed with 4% paraformaldehyde at 4°C overnight, washed 3 times with PBS and dehydrated in 30% sucrose in PBS for 3 days. Then, the brain was cut into 6 pieces at the positions of 5, 2, -1.5, -5.5 and -9 mm relative to the bregma (Supplemental Figure 1). Each piece was individually embedded into OCT compound and frozen. Finally, 11 coronal slices of 100 µm were cut at 7, 6.5, 5, 4, 2, 1, -1.5, -2.5, -5.5, -6.5 and -9 mm relative to the bregma to examine the tracer distribution using a fluorescent microscope (AX10, Zeiss, Oberkochen, Germany). Separately, 3 coronal slices of 10 µm were cut at 4, -2.5 and -6.5 mm relative to the bregma, representing the anterior, middle and posterior regions, respectively, to examine microglia activation.

Frozen blocks of the dCLN were prepared in the same manner as the brain. Two slices of 100 µm were cut 400 µm apart for each dCLN to examine the tracer distribution. All images of the brain and dCLN were quantified by ImageJ software (NIH, Bethesda, MD) with an identical threshold value.

For the meninges, the skull was fixed with 4% paraformaldehyde for 6 hours and kept in PBS overnight. The meninges were then detached from the skull, mounted on slides and kept at 4°C for 1 hour to let them dry and attached to the slides. The tracer distribution was assessed using a fluorescent microscope (AX10, Zeiss). The cut points of the anterior, middle and posterior regions were at 7, 1.5, -4 and -9.5 mm relative to the bregma, respectively (Supplemental Figure 1). Each part was 5.5 mm in width.

For biochemical analysis, the brain was cut into the anterior, middle and posterior regions at 7, 1.5, -4 and -9.5 mm relative to the bregma immediately after isolation. The cerebral cortex was dissected from each region, immersed into liquid nitrogen and preserved in -80°C until used for mRNA isolation and analysis of key metabolites (ammonia, glutamate and glutamine). The meninges were immersed into liquid nitrogen without segmentation and kept in -80°C for mRNA isolation.

***Immunofluorescence analysis of the brain and meninges***

A frozen brain was sectioned at 10 μm to examine microglia activation as mentioned above. Sections were incubated in blocking buffer (5% donkey serum and 0.3% Triton in PBS) for 1 hour and incubated with a primary antibody (rabbit anti-Iba1, 1:100, 019-19741, Wako Chemicals USA, Richmond, VA) at 4°C overnight. After washing 3 times with PBS, they were incubated with a secondary antibody (donkey anti-rabbit conjugated with Alexa Flour 546, 1:300, Invitrogen, Carlsbad, CA) in PBS containing 5% donkey serum and 1% bovine serum albumin for 1 hour. After washing 3 times with PBS, tissues were mounted with mounting media containing DAPI (Fluoroshield, Sigma-Aldrich, St. Louis, MO). Slides were evaluated for microglia morphology using a fluorescent microscope (AX10, Zeiss).

The meninges mounted on slides and kept at 4°C for 1 hour were incubated in blocking buffer overnight and then in primary antibodies (rabbit anti-Prox1, 1:500, DP3516P, Acris Antibodies GmbH, Hiddenhausen, Germany; mouse anti-CD31, 1:100, BD Pharmingen, San Diego, CA; rabbit anti-Iba1, 1:100, Wako Chemicals USA; mouse anti-CD68, 1:200, MCA341GA, AbD Serotec, Oxford, UK) at 4°C for 2 nights. After washing 3 times with PBS, the meninges were incubated with a secondary antibody (donkey anti-rabbit conjugated with Alexa Flour 546 or donkey anti-mouse conjugated with Alexa Flour 488, 1:300, Invitrogen). After washing 3 times with PBS, the meninges were mounted with Fluoroshield with DAPI (Sigma-Aldrich) and assessed for lymphangiogenesis and tracer uptake within cells using a fluorescent microscope (AX10, Zeiss).

***Quantification of meningeal lymphangiogenesis***

In the meninges, the area surrounded by the cerebrum olfactory bulb fissure (COF), superior sagittal sinus (SSS) and transverse sinus (TS) was analyzed by ImageJ. Prox1 positive nuclei were connected using the paintbrush tool in ImageJ, which represented lymphatic vessels. The total length of lymphatic vessels was measured. Meningeal lymphangiogenesis was expressed as the ratio of the total lymphatic vessel length to the meningeal area analyzed.

***Measurement of key metabolites***

Ammonia, glutamate and glutamine levels were evaluated in the anterior, middle and posterior regions of the cerebral cortex, using their respective assay kits (ammonia, ab83360, Abcam, Cambridge, UK; glutamate, ab138883; and glutamine, ab197011), according to the manufacturer's instructions. In brief, each part of the cortex was crushed into powder while frozen in the presence of liquid nitrogen. Powered tissues were homogenized with specific assay buffers provided by the kits. Total protein levels were determined by a protein assay (Bio-Rad Laboratories, Hercules, CA) prior to measurement of the metabolites. The results were expressed by metabolite amounts/ amount of total protein.

***Rotarod performance test***

Rotamex-5 (Columbus Instruments, Columbus, OH) was used for testing the ability of rats to stay on a rotating rod[24]. The test was performed one day before sample collection at the end of 4-week BDL. Rotating speeds were increased from 4 to 36 rpm over 300 seconds. The length of time for which a rat was able to stay on the rotating rod was recorded. The maximum cut-off time was 300 seconds. The average time of 5 consecutive tests represented motor function of a rat.

***RNA-seq library preparation and sequencing***

Total RNA was isolated and purified using the RNeasy Mini Kit (Qiagen) according to the manufacturer's instructions. RNA quality and integrity were verified by micro-volume spectrophotometry (NanoDrop 2000, Thermo Fisher Scientific) and by on-chip electrophoresis (2100 Bioanalyzer, Agilent Technologies, Santa Clara, CA), respectively. An $OD_{260/280}$ ratio of 1.8-2.0 and a RNA integrity number (RIN) of 8 or higher were considered to qualify intact RNA for further processing. Aliquots of the total RNA were sent to the Yale Center for Genomic Analysis for library preparation.

Library preparation was performed using the KAPA mRNA HyperPrep Kit (Roche Sequencing Solutions, Pleasanton, CA) according to the manufacturer's instructions. Briefly, mRNA was purified from 200ng of the total RNA with oligo dT beads and sheared into short segments through incubation at 94°C in the presence of magnesium ($Mg^{2+}$). Subsequently, the first-strand cDNA was synthesized by random primers. Second-strand synthesis and A-tailing were performed with dUTP to generate strand-specific sequencing libraries. Libraries were then sequenced using the NovaSeq 6000 System (Illumina) with parameters set for high output, paired-end and 100-bp sequencing. Samples were loaded onto the NovaSeq flow cell at a concentration that could yield 25 million passing filter clusters per sample.
